# PARP1 inhibition regulates tumor progression through modulation of RhoGDIα and vimentin in triple negative breast cancer

**DOI:** 10.64898/2026.07.18.739208

**Authors:** Jyotika Rajawat, Nidhi Shukla, Akanchha Shukla, Manohar Singh, Aijaz Ahmad John, Divya Singh, Mandira Sharma, Durga Prasad Mishra

**Affiliations:** CSIR-Central Drug Research Institute (CDRI); Lucknow Cancer Institute

**Keywords:** Poly(ADP-ribose) polymerase1, RhoGDIα, Vimentin, metastasis, breast cancer, BRCA

## Abstract

**Background and Purpose:** PARP inhibitors have been evaluated in clinical trials for several cancers and Olaparib is FDA approved for treating BRCA deficient ovarian cancer. Numerous reports have suggested Poly(ADP-ribose) polymerase1(PARP1) overexpression in a variety of cancers including breast carcinomas and proposed the role of PARP1 in metastasis. However, the mechanism of PARP1 in regulating metastatic process in BRCA proficient and deficient TNBC is not studied thoroughly. In this study, we propose that PARP1 mediated breast carcinoma progression is gene transcription mediated, where it regulates several steps of pro-metastasis.

**Experimental Approach:** PARP inhibitor’s effect on metastasis was monitored by migration and invasion assay, modulation in protein expression was assessed by proteomic analysis and further confirmed by immunoblotting. Chromatin immunoprecipitation was performed to study the transcriptional role of PARP1. Ectopic expression and siRhoGDIα, and immunofluorescence assessed the cytoskeleton changes. PARP inhibitor was administered in xenograft mice to study metastasis. Immunohistochemical analysis was done on patient and mice tissues.

**Key results:** Breast cancer cells exhibited reduced migration and invasion due to PARP1 inhibition. PARP1 regulates expression of vimentin and RhoGDIα and hence cytoskeletal rearrangement causing a change in migrating potential of a cell. Metastasis in mice was reduced upon PARP inhibition. PARP1 was identified to be a novel transcriptional regulator of RhoGDIα. Furthermore, RhoGDIα ectopic expression substantiated the PARP inhibitor effects, suggesting the PARP inhibitor downstream signaling to be mediated through RhoGDIα.

**Conclusions and Implications:** We identified a novel aspect of PARP1 as promoter of metastasis via transcriptional regulation of RhoGDIα. Assessing RhoGDIα levels in TNBC patients might be useful to predict sensitivity to PARP inhibitors.

## Introduction

Poly (ADP-ribose) polymerase1 (PARP1) is one of the first respondent protein in DNA damage response (DDR), where it recruits multiple DNA repair proteins on the damaged DNA site and eases the process of repair (Wei & Yu, 2016). Being a part of repair protein in the DDR pathway, its inhibition along with the DNA damage inducing agent has revolutionized the therapeutic approach in cancer (Brown et al., 2017). The synthetic lethality approach, using PARP inhibitor along with the DNA damaging agent has been evaluated in clinical trials for BRCA deficient cancers (Farmer et al., 2005). PARP inhibitors, Olaparib has been approved by FDA for treatment of BRCA deficient ovarian cancer. Recent phase I trial of Olaparib in combination with capivasertib (AKT inhibitor) exhibited a synergistic effect on BRCA1/2 deficient and proficient tumors (Yap et al., 2020). Further research and clinical trials are underway for current generation PARP inhibitors in solid tumors and leukemia irrespective of BRCA defect.

Pleiotropic role of PARP1 as transcription regulator and in epigenetics have signified its role in cancer progression (Rajawat et al., 2017). Contemporary findings have highlighted the role of PARP1 in regulation of several transcription factors, whereby it can up or down regulate expression of genes (Hassa et al., 2003). Numerous reports have suggested overexpression of PARP1 in a variety of cancers, including leukemia (Gil-Kulik et al., 2020; Pashaiefar et al., 2018), sarcomas (Kim et al., 2016) and carcinomas of breast (Siraj et al., 2018; Stanley et al., 2015), ovary (Marques et al., 2015), glioblastoma (Galia et al., 2012), liver (Guillot et al., 2014) and colon (Dziaman et al., 2014). Thus, PARP1 is an important factor that seems to be involved in controlling various changes during tumor progression and metastasis (Bai & Virag, 2012; Brenner et al., 2011; Curtin & Szabo, 2013; Feng et al., 2015; Schiewer & Knudsen, 2014). Recent studies have demonstrated that PARP1 promotes metastatic phenotypes through interactions with transcription factors such as ZEB1, NF-κB, and Smad4 (Kumar et al., 2021; Lin et al., 2024; Schacke et al., 2019). Pharmacological inhibition of PARP1 has been shown to suppress epithelial to mesenchymal transition (EMT), reduce migratory and invasive capabilities, and decrease distant metastatic spread in several cancer models (Beniey et al., 2023; Choi et al., 2016a; Frederick et al., 2024). However, the mechanism of PARP1 in regulating metastatic process in BRCA proficient and deficient triple negative breast cancer (TNBC) at molecular level is still not completely elucidated.TNBC is highly aggressive and fatal cancer associated with poor outcome due to its characteristic feature of distant metastasis. Chance of recurrence in TNBC is high due to its heterogeneous nature with limited therapeutic options, furthermore, resistance to chemotherapy. Metastasis in TNBC is a complex multistep process and is poorly understood.

Understanding the cellular and molecular changes associated with cancer progression and metastasis is of prime importance. Cancer progression is a multistep process which also involves change in proteins responsible for cell to cell contact (Kaminska et al., 2015). Proteomic analyses have revealed that PARP inhibition alters the expression of metastasis-associated proteins, particularly those involved in cytoskeletal organization and cell–cell adhesion, highlighting a broader role of PARP1 in cancer dissemination (Rajawat et al., 2023). Tight junction and adhesion proteins together assist in cell-cell adhesion. Cytoskeleton remodelling and intermediate filament rearrangement is a prerequisite phenomenon during cell migration and EMT. Cytoskeleton protein RhoGDIα is a regulatory protein that have significant role in microtubule dynamics, cell polarity and cytoskeleton regulation. RhoGDIα dictates the spatial and temporal pattern of RhoGTPases activation. Altered expression of RhoGDIα has an impact on activity and downstream signaling of RhoGTPases and hence on cell motility (Bozza et al., 2015). RhoGDIα expression varies in several tumors and loss of RhoGDIα has been proposed to be a prognostic factor in hepatocellular carcinoma(Li et al., 2013). RhoGDIα downregulation promotes cancer cell growth and stem cell maintenance (Wu et al., 2016; Xiao et al., 2014; Zhu et al., 2017; Zhu et al., 2012). Contrarily, RhoGDIα is known to be involved in tumor cell apoptosis as well as migration of a tumor (Zhang et al., 2005). Vimentin is one of the key intermediate filaments mostly expressed in mesenchymal cells and hence is one of the indicators of cancer progression (Vasko et al., 2007). Vimentin exhibits or modulates cell-cell contact and migration of cell either directly or indirectly(Singh et al., 2003). PARP1 has been seen to play a role in the increased transcription of vimentin in lung cancer cells through the activation of its promoter (Chu et al., 2007).

Several studies recently have proposed potential benefits of PARP inhibitors in cancers with no defects in repair pathway. However, molecular mechanisms of other functions of PARP1 particularly in context to cytoskeletal proteins are yet to be characterized. The rationale of the present study focuses on delineating the molecular link involved in PARP1 mediated breast cancer progression. The mechanistic study will help in elucidating the novel regulatory functions of PARP1 and identifying patient population for selective treatment of cancer depending on different stages and response to PARP inhibitors.

## Materials & Methods

### Cells and Cell culture

Breast carcinoma cell lines MDA-MB-231 and HCC1937 were maintained in a phenol free RPMI, and MCF7 in MEM supplemented with 10% fetal bovine serum and 1X antibiotic/antimycotic solution (Himedia, India) in an incubator with 5% CO_2_ at 37°C.

### Plasmids and siRNA transfection

si*RhoGDIα* was purchased from Santa cruz Biotechnology and RhoGDIα overexpression vector from GenScript (NJ, USA). Transient transfection was performed using Xfect transfection reagent (Clontech) according to manufacturer’s protocol.

### Clonogenic assay

To evaluate cell colony forming ability, cells were plated in 12 well plates at a density of 500 cells/ml.Next day, cells were treated with PARP inhibitor, Olaparib (Cayman chemicals) for 24 hr. Medium was removed and fresh complete medium was added and incubated for one week.Colonies were stained with Coomassie blue and counted. Percentage of colonies arising from surviving cells was calculated relative to colonies arising from control cells.

### Cell migration assay

Cells were plated at high density to reach 90-100% confluence 24 hours later, a scratch was made in the middle of the dish with a sterile 200µL micropipette tip, media removed, washed with PBS and fresh media replenished, with addition of Olaparib in treated cells. Images were taken after 0-24 hr at a 20X magnification. Area of wound was quantified by ImageJ software and the percent of wound closure was determined.

### Invasion assay

Invasion assay was performed using matrigel coated transwell inserts, where lower chamber was filled with serum containing media while the cell solution in serum free media containing Olaparib was added on the top chamber. Cells were incubated for 24-30 hrs and then processed for staining as described by Yang et al., 2014. The migrated cells invaded to the other side of the membrane were viewed under inverted microscope at 20X magnification.

### Cell adhesion assay

For adhesion assay 25000 cells were seeded on fibronectin coated 96 well plate. Cells were allowed to adhere for 3 hr and then washed with 1X PBS. Cells were fixed with 4%paraformaldehyde for 10 min, washed and then stained with crystal violet at room temperature for 10 min. Cells were washed 5 times with water, followed by extraction of dye with 0.2% Triton X-100. Optical density was measured at 595nm and percentage of adhesion was normalized to control cells.

### Western Blot analysis

Total cell lysates were prepared in RIPA buffer supplemented with protease inhibitors (Sigma-Aldrich). Lysates were kept on ice and then centrifuged at 14,000 g for 30 min at 4°C. Supernatants were collected and quantified by Bradford protein assay reagent (Sigma). 40μg of proteins were loaded on SDS-PAGE and electroblotted onto nitrocellulose membrane (Millipore). Immunoblots probed with the specific antibodies were developed using ECL Plus chemiluminescence reagent (Millipore). We used EMT sampler kit (Cell Signaling Technology) to identify the change in expression of PARP1 targeted EMT proteins. Other antibodies used were: PARP1 (abcam), RhoGDIα (abcam) and Actin (Santa Cruz Biotechnology).

### Immunoprecipitation

Cells were lysed with RIPA lysis buffer containing protease inhibitor cocktail, incubated for 30 min in ice and lysate was spun at 14,000 g for 30 min. Protein was estimated by Bradford assay and 1mg/ml protein was used. The supernatants were incubated with anti-PARP1 antibody for overnight at 4°C. The beads were washed with IP buffer, then added in the lysate IP complex and incubated for 2 hrs. The immunoprecipitates were washed three times with buffer and beads were then eluted with sample buffer and heated for 10 min at 70°C. The supernatant was collected, beads were removed, protein loaded on the gel and analyzed by western.

### Chromatin immunoprecipitation (ChIP)

Chromatin was isolated from cells using ChIP kit (CST) and further purified and eluted as per the protocol from the manufacturer. Immunoprecipitation was done with the isolated chromatin using PARP1 antibody and further processed as given in the protocol. Chromatin was extracted from IP beads, purified and quantified and further PCR was carried out using gene specific primers for ChIP.

### *In Vivo* Tumor Generation and metastasis

4T1 syngenic breast cancer metastasis model was done as described previously (Thakur et al., 2015). 5 female balb/c mice were taken in each group. Control group was orally administered with vehicle (10% DMSO in PBS) and treatment group with 25mg/kg body weight of olaparibdaily for 3 weeks. MicroCT was performed to analyse bone metastasis.

### Immunohistochemistry

Paraffin-embedded paraformaldehyde blocked tissue samples (patients and control) were stained for the expression of PARP1, vimentin and RhoGDIα as described previously (Galia et al., 2012). Antibody binding was visualized with goat anti-rabbit or anti-mouse antibody coupled to horseradish peroxidase (Santa Cruz) and 3,3’-diaminobenzidine. Counterstaining was performed with Harris’ modified hematoxylin (Sigma, USA).

### Statistical analysis

Graphpad prism software was used to perform statistical analyses. One-way Anova or students t-test was applied for comparing the experimental data.

## Results

### Lower PARP1 expression correlates with metastasis free survival in triple negative breast cancer (TNBC) patients

To assess the potential role of PARP1 in triple negative breast cancer progression, we explored the correlation between the expression of PARP1 and its association with disease free survival in patients using *in silico* analysis in triple negative breast cancer patients. Analysis of the data of TNBC patients in The Cancer Genome Atlas (TCGA) dataset and other cohort studies indicated a higher expression of PARP1 (Figure 1A). We then investigated the association of PARP1 levels and overall survival in triple negative breast cancer patients from the data available on Geo database. The analysis revealed that the elevated PARP1 levels were associated with the reduced overall patient survival (Figure 1B). This association of PARP1 with poor overall survival was independent of age, gender and cancer stage. We also explored the involvement of PARP1 expression in metastasis free survival in triple negative breast cancer to assess its role with metastasis from the reported dataset on the Geo database. An association between higher PARP1 expression and reduced metastasis free survival as well as reduced lung metastasis free survival was observed in two different cohorts GSE58812 and GSE5327 respectively (Figures 1 C and D). As the higher mortality rate in triple negative breast cancer patients is due to relapse of tumor, we further investigated the association of PARP1 expression with relapse free survival. Analysis of the Cohort study GSE2607 indicated the significantly poor relapse free survival in patients with higher PARP1 expression (Figure 1E). A meta-analysis study reported that elevated PARP levels correlates with worse prognosis in early stage breast cancer (Qiao et al., 2017, PloS One). Collectively, these analyses of the cohort studies indicated that the elevated PARP1 expression is associated with poor overall, metastasis free and relapse free survival thereby underscoring its potential role in breast cancer progression and metastasis.

**Figure 1.**
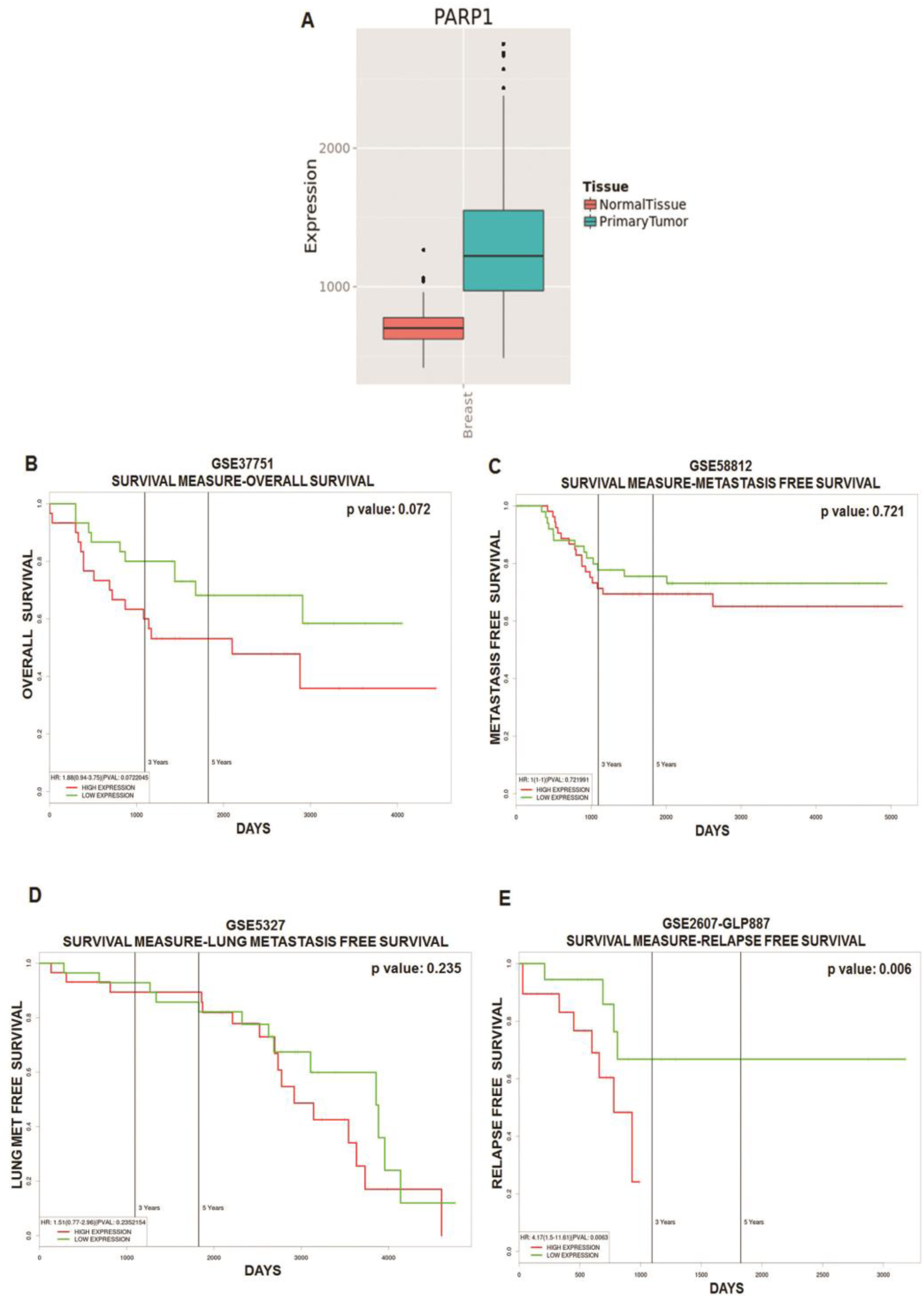
PARP1 association with metastasis free survival in triple negative breast cancer patients. (A) Elevated expression of PARP1 in breast cancer tissue compared to normal breast tissue. The data of breast cancer from TCGA dataset indicated significant higher expression of PARP1 in breast cancer tissue. (A) n =108; (B-E) PARP1 expression and survival relationship in breast cancer patients. (B) n=94 (p value=0.072); (C) n=107 (p value-0.721); (D) n=58 (p value= 0.235); (E) n=47,38 (p value= 0.006). Data was gained from Geo database, and then Kaplan Meier survival curve was generated using online software known as PROGgene. Survival curve exhibited the PARP1 association with overall survival, metastasis free survival, lung metastasis free survival and relapse free survival.

### PARP1 regulates proliferation, migration, invasion and clonogenicity in breast cancer cells

The *in silico* analysis indicated a role for PARP1 in triple negative breast cancers (Figure 1). Therefore, we next assessed the association of PARP1 with cancer hallmarks in triple negative breast cancer cell lines. We carried out these studies in BRCA positive (MDA-MB-231 and MCF7) and BRCA negative (HCC1937) cells. For these studies we usedolaparib, a competitive inhibitor of PARP1. First, the effect ofthe PARP inhibitor olaparibon proliferation of breast cancer cells was studied using MTT assay. The results indicated a significant difference in cell survival at 48 hrs post treatment. Approximately 50% cell death was observed in the olaparib (80µM) treated cells after 48 hrs of treatment in all breast cancer cell lines (Figure 2A). Further experiments were carried out at a sub lethal dose (10µM).

**Figure 2.**
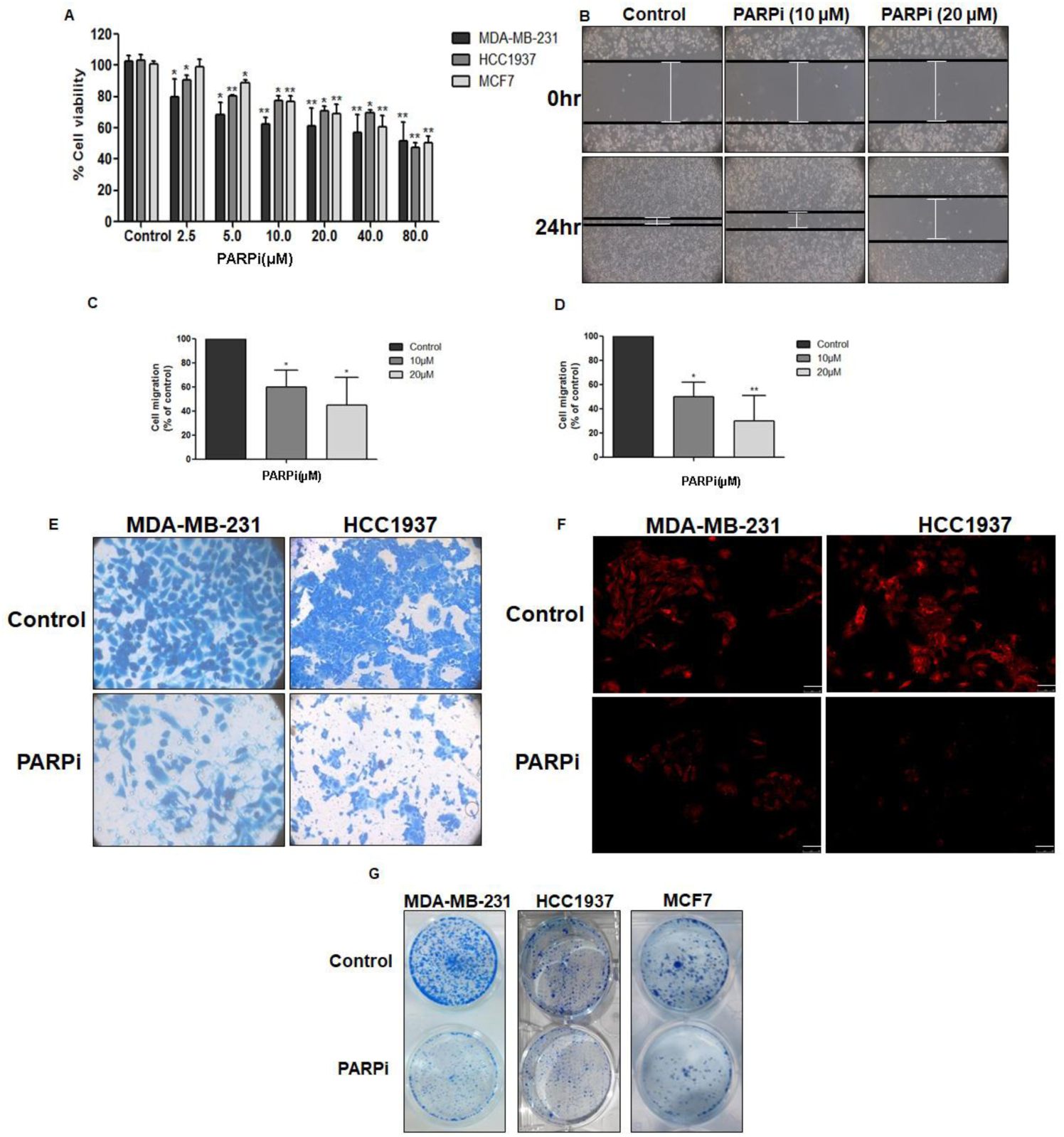
PARP1 inhibition suppresses cell growth, migration and invasion in breast cancer cells. (A) MTT assay after 48 hrs of olaparib treatment (B) Migration ability was monitored by scratch assay and images were taken after 24 hrs in light microscope at 10X magnification (C, D) cell migration as % of control in MDA-MB-231 and HCC1937 cells respectively (E) Cell invasion examined by matrigel invasion chamber (F) Phalloidin staining to monitor the changes in cytoskeleton arrangement after PARP inhibition in MDA-MB-231 and MCF7 cells. Images taken in 60X magnification and scale bar is 100µm. (G)Colony formation in PARP1 inhibited breast cancer cells All data represent mean ± SD from three independent experiments and p-values were calculated by students t-test, *<0.05, **<0.01.

Metastatic potential is the significant feature of breast cancer cells, and migration and invasion are key processes of the metastatic cascade (van Zijl et al., 2011). The highly metastatic TNBC cell line MDA-MB-231 was used to investigate the effect of PARP inhibition on migration and invasion potential of breast cancer cells by the *in vitro* wound healing and matrigel invasion assays respectively. Olaparib significantly inhibited the migratory and invasive ability of breast cancer cells (MDA-MB-231 and HCC1937) at a sub lethal dose. Wound healing assay exhibited 60% and 45% of migration with 10 and 20µM olaparib treatment respectively in MDA-MB-231 cells compared to that of the control cells (Figures 2B & 2C). Moreover, HCC1937 cell migration was also inhibited after 24 hrs of olaparib treatment, where 50% and 30% migration were observed compared to that of the control cell (Figure 2D). Similarly, olaparib treatment significantly inhibited invasion of cells by 72% and 68% in MDA-MB-231 and HCC1937 cells respectively (Figure 2E). Next, we investigated the cytoskeleton rearrangement upon olaparib treatment by analysing F-actin using phalloidin staining. Interestingly, we observed that cell polarity was affected after treatment with PARP inhibitor (Figure 2F).

We finally performed the clonogenic assay as a surrogate to the *in vivo* animal models and found that the survival fraction of PARP inhibitor treatment inhibited clonogenicity in the breast cancer cells (Figure 2G). Collectively, these results demonstrated the involvement of PARP1 in regulation of cancer hallmarks in triple negative breast cancer cells.

### PARP1 regulates metastasis by targeting cytoskeletal and adhesion proteins

To characterize the mechanism of migration and invasion regulated by PARP1, we performed a proteomic analysis of MDA-MB-231 cells in the presence and absence of olaparib. The expression levels of a number of proteins were altered following PARP1 inhibition, as detected by the 2D gel electrophoresis (Figure 3A) and mass spectrometry analysis. A two-fold down-regulation of vimentin (a class III intermediary filament), 2-phosphopyruvate-hydratase alpha-enolase, glyceraldehyde-3-phosphate dehydrogenase, 14-3-3 protein epsilon was observed following PARP inhibition. An increase in the expression of the protein disulfide isomerase PDIA6, proteasome subunit alpha type-3 isoform 1, Rho GDP-dissociation inhibitor 1 isoform-α (RhoGDIα) was also observed after PARP1 inhibition (Figure 3A and Supplementary Table S1). Among these proteins, we found RhoGDIα and vimentin to be involved in cell migration critical to the metastatic process.

**Figure 3.**
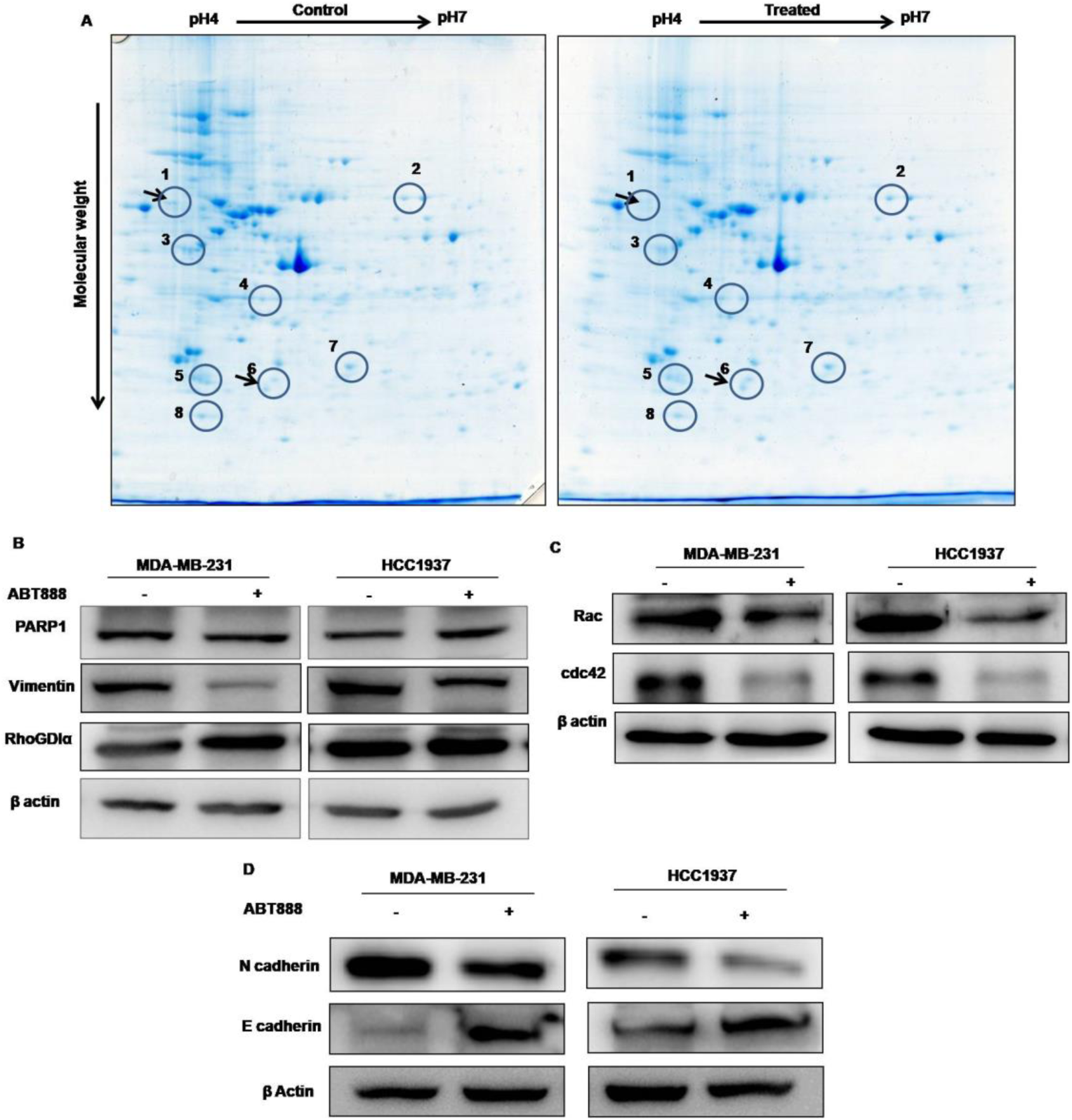
Proteomic profiling of untreated and PARP1 inhibitor treated MDA-MB-231 cells. (A) Change in proteomic profile of MDA-MB-231 cells upon PARP inhibition by 2D gel electrophoresis and mass spectrometry. Arrow indicates spot of vimentin (1) and RhoGDIα (6) protein which were downregulated and upregulated respectively after olaparib treatment. Differentially expressed spots were excised and analyzed by MS/MS (B) Western analysis of PARP1, vimentin and RhoGDIα proteins (C) Western analysis of cytoskeletal proteins (Rac & cdc42) in MDA-MB-231 and HCC1937 cells (D) Western analysis of EMT proteins (N cadherin and E cadherin). Data is a representation of three independent experiments.

The expression levels of these differentially expressed proteins in 2-D/MS were then analysed by immunoblotting. Our results confirmed reduced expression of vimentin in both MDA-MB-231 and HCC1937 cells following PARP inhibition. Conversely, cytoskeletal protein RhoGDIα was increased upon PARP1 inhibition in MDA-MB-231 cells (Figure 3B). Hence, we also checked the expression of proteins involved in RhoGDIα signaling which indicated reduction in the expression of Rac and Cdc42 (Figure 3C). RhoGDIα levels were not noticeably changed in HCC1937 cells but decreased expression of Rac and Cdc42 were observed (Figure 3C). RhoGDIα depletion also inhibits migration and invasion due to misfolding of Rho proteins and further degradation (Boulter et al., 2010), or hyperactivation of RhoGTPases and hence their uneven distribution in cell leads to inhibition of cellular migration (Reyes et al., 2013).

Most cells acquire migratory phenotype after undergoing epithelial to mesenchymal transition (EMT). Hence, we next checked the expression of selected proteins known to be involved in EMT. The results showed that the expression of mesenchymal protein N-cadherin, was diminished while of epithelial protein E-cadherin was increased in both MDA-MB-231 and HCC1937 cells, indicating a disruption of EMT in the absence of PARP1 activation (Figure 3D). Collectively, the data suggests that PARP1 activity regulates the expression of these cytoskeletal and adhesion proteins, thereby regulating metastasis.

### PARP1 transcriptionally regulates RhoGDIα and vimentin

PARP1 is reported to modulate transcription of various genes through direct or indirect interaction with promoter or transcription factors (Kraus, 2008). Our results suggested the change in protein expression of vimentin and RhoGDIα upon PARP inhibition. To investigate whether PARP1 acts as a transcriptional regulator of these two proteins, we first performed immunoprecipitation followed by MS/MS analysis. The results revealed the presence of vimentin in the PARP1 IP complex (Figure 4A). Further ChIP analyses confirmed direct interaction of PARP1 with RhoGDIα and vimentin (Figures 4C-D). The result showed the binding of PARP1 to the promoter of RhoGDIα and vimentin, and thus regulating their expression in breast cancer cells.

**Figure 4.**
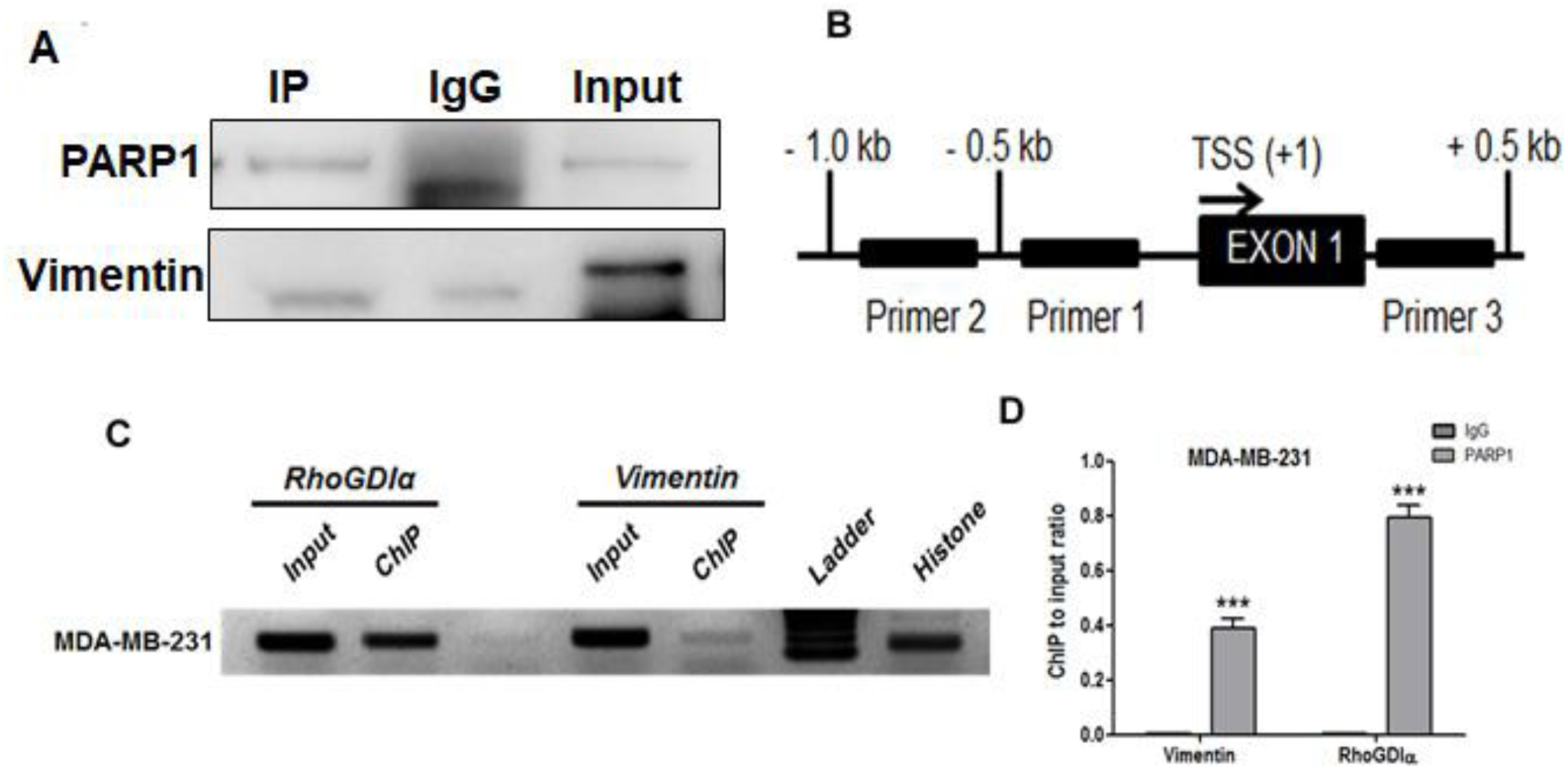
Transcriptional regulation of RhoGDIα and Vimentin by PARP1. (A) Immunoprecipitation with PARP1 antibody in MDA-MB-231 cells to detect PARP1 interacting partners. Immunoblot with vimentin antibody after immunoprecipitation (B) ChIP primers designing strategy, where three primer set were designed for each gene. Positive binding was seen with only one primer located upstream to exon 1 (C) Chromatin immunoprecipitation performed using PARP1 antibody and IgG; suggested PARP1 as direct regulator of vimentin and RhoGDIα. (D) Quantitation of enrichment of PARP1 as percent input. Data is a representation of three independent experiments; p-values were calculated by students t-test, ***<0.001.

### RhoGDIα expression inhibits migration and invasion in breast cancer cells

To investigate the function of RhoGDIα as mediator of PARP1 during breast cancer metastasis we employed loss and gain function strategy. MDA-MB-231 cells were transfected with RhoGDIα expression vector, empty vector, as well as siRNA. Cell growth was affected after transfection, as observed (Figure 5A). Clonogenic activity was checked after transfection and was found to be reduced upon expression of RhoGDIα expression vector, while no significant effect was observed with siRhoGDIα (Figure 5B). We next used matrigel invasion chambers to further monitor the effect of RhoGDIα expression on invasion, whereby reduction in invasion capacity was observed compared to that of the controls (Figure 5C). Cell adhesion was also significantly reduced by 20% and 30% upon PARP inhibition and RhoGDIα expression respectively (Figure 5D). For migration assay, transfected cells were scraped at confluence and monitored for migration (wound closure) after 24 hrs. Intriguingly, we observed reduction in migration of MDA-MB-231 cells transfected with RhoGDIα vector (Figure 5E). Similar results were observed with olaparib, thus supporting the involvement of RhoGDIα in migration and invasion of breast cancer cells. Rho family members are known to be involved in maintaining the cytoskeletal structures and actin polymerization (Yu et al., 2012, JBC). We therefore subsequently studied the effect of RhoGDIα on cytoskeletal rearrangement by using phalloidin staining. Interestingly, RhoGDIα expression modified the actin rearrangement similar to that of PARP inhibition while, no change was observed with the genomic inhibition of RhoGDIα (Figure 5F). Collectively, these data suggest that PARP1 regulated tumor progression is mediated through RhoGDIα.

**Figure 5.**
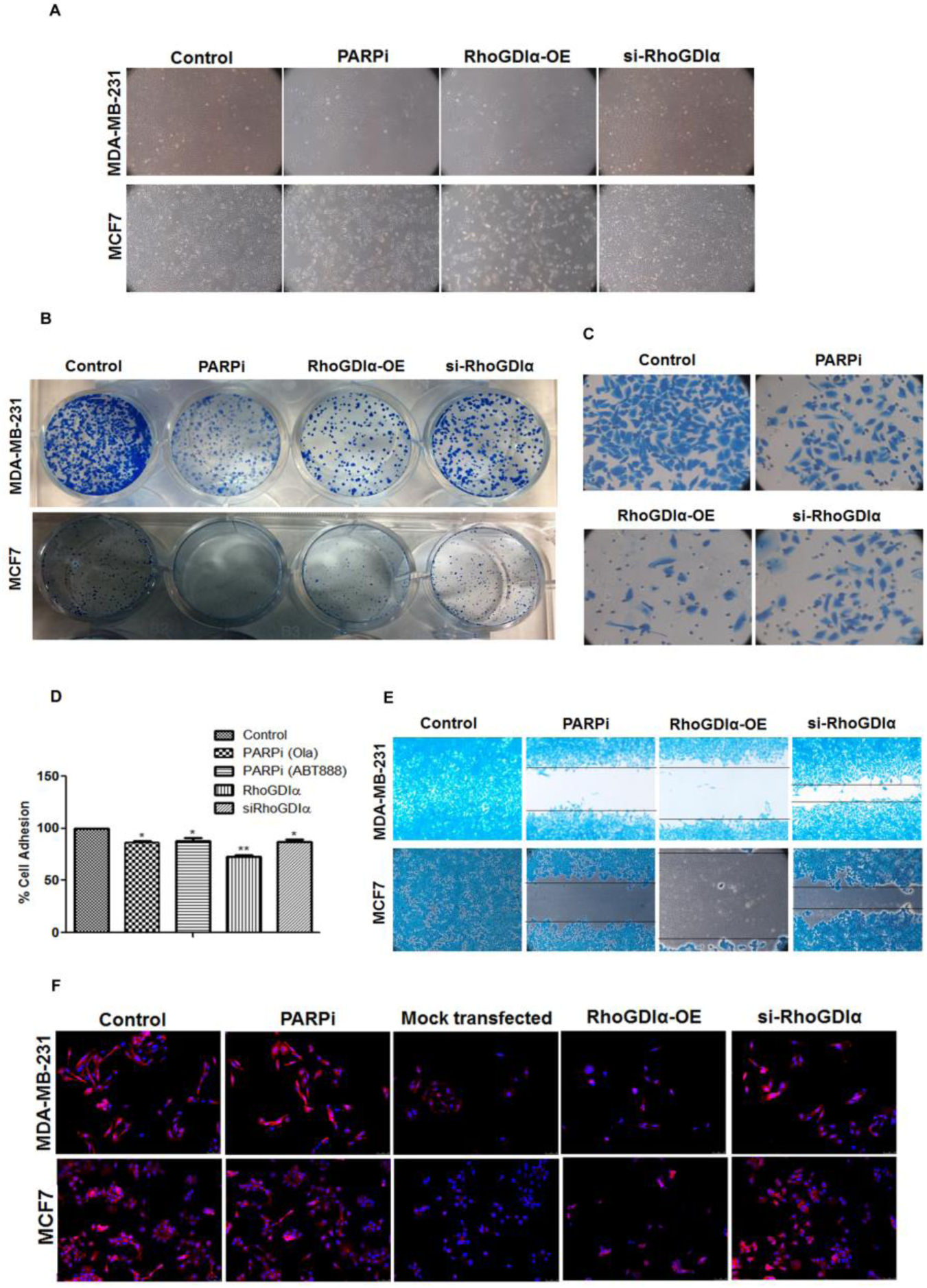
RhoGDIα expression correlates with PARP inhibition. (A) Cell growth monitored after 24 hrs of transfection (B) Clonogenic assay, where cells were seeded in 12-well after 24 hrs of transfection (C) Matrigel invasion assay 24 hrs post transfection (D) Cell adhesion assay using fibronectin coated wells to assess the adhesion capability of breast cancer cells (E) Cell migration potential in MDA-MB-231 cells, representative images of migrating cells stained with coomassie blue (F) Phalloidin staining for cytoskeletal rearrangement. Images were taken in 60X magnification and scale bar is 100µm.Data represent mean ± SD of three independent experiments.

### PARP inhibition impairs tumor growth *in vivo*

To investigate the tumor suppressive capacity of PARP1 inhibitor *in vivo*, the 4T1 breast cancer cells (2×10^6^ cells) were subcutaneously injected into mammary fat pads of 4-week old Balb/c mice. Tumor growth was monitored for one week and animals were grouped (5 animals/group), followed by oral administration with PARP inhibitor, olaparib (25mg/kg daily for 3 weeks) (Drew, 2015). Inhibitory effect of PARP suppression on tumor growth was evident from day 7 of treatment and consistent reduction in tumor size was observed during the sacrifice of mice. Animal body weight was recorded during the course of experiment and no prominent change was observed (Figure 6A). Tumor volume was measured *ex vivo* which further confirmed the reduction in tumor size (Figures 6B and D). Reduction in tumor volume (75% reduction) along with decrease in tumor weight was observed in PARP inhibitor administered mice (Figure 6C). No apparent toxicity was observed in liver and lung by PARP inhibition as assessed by haematoxylin and eosin staining (Figure 6E). We also determined the effect of PARP inhibition on metastasis and distant metastasis. Intriguingly, higher metastatic lesions were observed in lung and liver of control group, while no lesions were noted in the distant organs in PARP inhibitor treated mice. PARP inhibition reduced the size and number of lung and liver metastatic nodules as confirmed in H & E staining also (Figures 6F and G). Furthermore, we evaluated the association of PARP1 with vimentin and RhoGDIα in tumor tissue by immunoblot analysis. These results suggested that PARP inhibition increases the expression of RhoGDIα and decreases the level of vimentin (Figure 6H), corroborating with our *in vitro* results. Majority of deaths in patients with breast cancer have been reported to occur via metastases to bone and lung. To investigate the functions of PARP inhibitor in bone metastasis, we performed microCT analysis. Interestingly bone metastasis was markedly reduced upon PARP inhibition as evident from (Figure 6I-N). MicroCT data analysis revealed that tumor bearing mice showed the formation of osteolytic lesions and loss of trabecular network in femur compared to sham group while these effects were reversed in PARP inhibitor treated group (Figure 6M and N). Further quantification of trabecular bone parameters i.e., BV/TV., bone volume/tissue volume ratio; Tb.Sp., trabecular separation; Tb.N., trabecular number; Tb.Th., trabecular thickness was also conducted. Quantification of the micro-CT data revealed that tumor bearing mice showed significantly decreased bone parameters, including BV/TV(Figure 6I), Tb.N (Figure 6J), and Tb.Th (Figure 6K), with a concomitant increase in Tb.Sp in femora (Figure 6L), while these parameters were reversed in PARP inhibitor treated group compared to sham group. Collectively, our data concluded that PARP inhibition significantly suppressed primary tumor growth and metastasis. Taken together, we demonstrated the involvement of PARP1 in regulating tumor growth and metastasis through vimentin and RhoGDIα.

**Figure 6.**
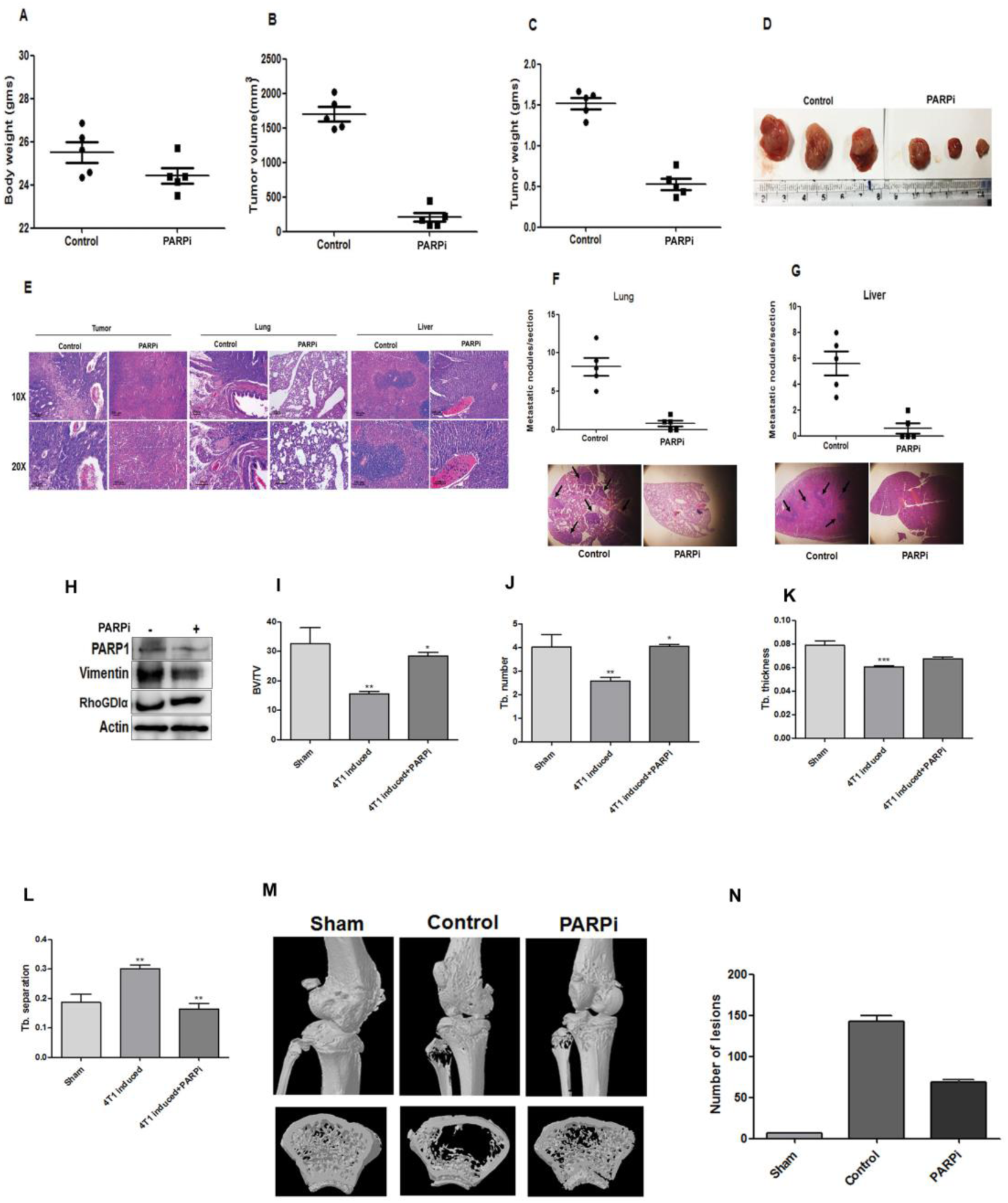
Therapeutic effect of PARP inhibitor on metastasis in 4T1 induced orthotopic tumor model of triple negative breast cancer. (A) Animal body weight before sacrifice (B) Tumor volume measured *ex vivo* using vernier calliper (C) Tumor weight taken after sacrifice (D) Tumor of vehicle and PARPi treated mice photographed after sacrifice (E) Representative images of H&E staining of tumor, lung and liver captured at 20X and 40X magnification (F, G) Quantification of metastatic nodules per section seen in lung and liver. Lower panel shows representative images of H&E staining captured at 4X magnification to show metastatic nodules in lung and liver of untreated mice. (H) Western analysis of PARP1, vimentin and RhoGDIα in tumor samples. (I-N) Bone metastasis was studied by microCT. Mice per group is 5.

### Immunohistochemical localization of PARP1 and its transcriptional targets in breast cancer tissues

Vimentin and RhoGDIα are cytoskeletal proteins involved in cytoskeleton rearrangement but their roles in breast cancer in the clinical setting are not clearly defined. Therefore, we analysed their status in triple negative breast cancer samples. The H&E staining revealed significant difference in morphology of breast tissue in control and TNBC patients (Figure 7A). To determine the level of these proteins and their correlation with PARP1, immunohistochemical analysis was carried out on triple negative breast tissue samples. The IHC analyses of PARP1 levels were carried out in breast cancer patients, whereby expression of PARP1 was predominantly nuclear (Figure 7B). Levels of vimentin were found to be increased in cytoplasmic region in tumor tissue samples (Figure 7B). However, RhoGDIα expression was not prominently observed in TNBC patients (Figure 7B). These results suggested clinical importance of PARP1, vimentin and RhoGDIα in biology of breast cancer progression and hence signified the importance of these proteins as biomarkers for PARP inhibition in chemotherapy.

**Figure 7.**
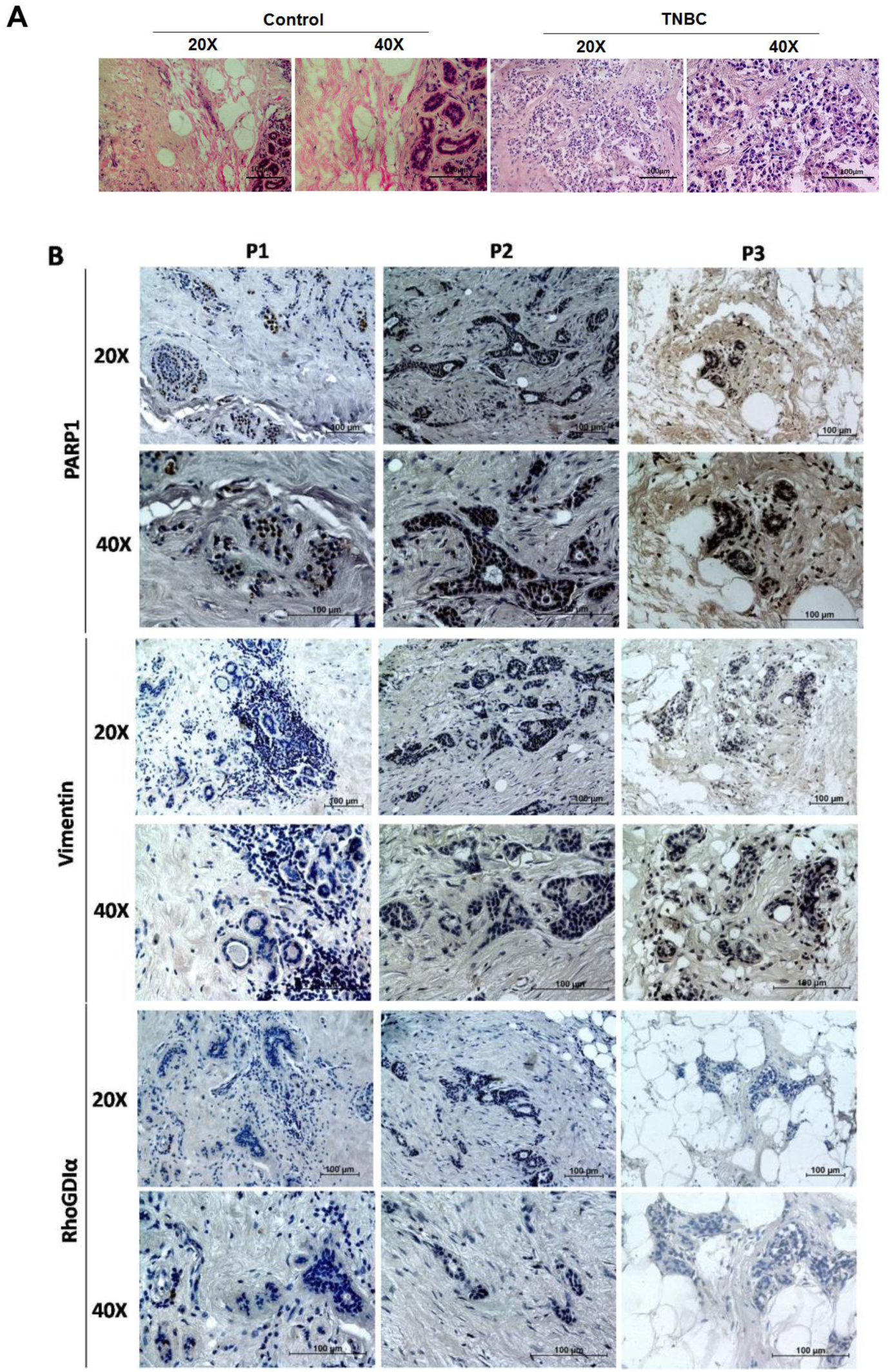
Clinical association of PARP1 with vimentin and RhoGDIα in triple negative breast cancer patients. (A) H&E of TNBC tissue and adjacent control (B) IHC of PARP1, vimentin and RhoGDIα in triple negative breast cancer patient tissues photographed in light microscope at 20X and 40X magnifications. Expression of PARP1 and vimentin was seen while expression of RhoGDIα was not prominent in TNBC patients (n= 10).

## Discussion

Determining the mechanism of action of PARP inhibitors, their adverse side effects and identifying the prognostic and treatment responsive biomarkers is imperative. Algorithm based studies have predicted gene based expression signature that will be useful for deciding synergistic therapies (McGrail et al., 2017). Such studies will elaborate on the future prospects of PARP inhibitors in clinical settings for cancer therapeutics. This study was carried out with two primary goals; firstly, to better understand the mechanistic role of PARP1 in metastasis in TNBC cells using PARP inhibitor olaparib and, secondly, to identify the signaling proteins regulated by PARP1 during metastasis. The current study reveals that PARP inhibitor olaparib inhibits tumor progression and metastasis by mediating changes in cytoskeletal and adhesion proteins. The key findings of this study are: (i) Olaparib treated TNBC cells exhibited reduced migration and invasion irrespective of BRCA deficiency (ii) Proteomic analysis after olaparib treatment revealed changes in cytoskeleton and adhesion proteins, particularly, RhoGDIα and vimentin in TNBC cells (iii) PARP1 is a novel transcriptional regulator of RhoGDIα and regulates cell migration and metastasis (iv) *in vivo* studies suggest that PARP inhibitor reduces tumor growth and also organ metastasis including bone metastasis.

PARP1 is no longer viewed only as repair protein; it plays multiple roles to perform and hence can be targeted per se depending on the function in specific cell or tissue type. Expression of PARP1 protein has been observed in several cancers including breast cancer and has been found to vary in subtypes (Bieche et al., 2013; Domagala et al., 2011; Green et al., 2015; Krukenberg et al., 2014). PARP1 behaves as a double edged sword, where PARP1 activation may lead to repair of damaged DNA, however, continued upregulation and activation may contribute to tumorigenesis (Swindall et al., 2013). Several studies recently have proposed potential benefits of PARP inhibitors in cancers with no defect in repair pathway. Recent reports have also proposed the role of PARP1 in metastasis process (Chen et al., 2017; Choi et al., 2016b). However, the mechanism of involvement of PARP1 in metastatic process in BRCA proficient and deficient TNBC is not studied thoroughly.

We analysed previously published data on metastasis free survival in relation to PARP1 expression and found that PARP1 level was inversely related to metastasis free survival and overall survival (Figure 1); indicating the role of PARP1 in metastasis. Aiming to investigate the possible cross talk between PARP1 and metastasis related EMT axis proteins in TNBC cells, firstly we monitored the PARP inhibitory effect on proliferative and migrating capability of TNBC cells using olaparib. Olaparib is a well reported PARP inhibitor that blocks PARP activity not only in cancer cells (Faraoni et al., 2015) but other organ cells too (Ahmad et al., 2018; Korkmaz-Icoz et al., 2018). Consistent with the previous findings, we also observed that PARP inhibition reduced the migration capacity and proliferation ability significantly but does not induce apoptosis. Our data suggest that PARP inhibition reduces the proliferation, migration and invasion in TNBC cells irrespective of BRCA gene (Figure 2). Barboro and colleagues have shown that PARP inhibitor treated prostate cancer cells could not migrate. Simultaneously, invasive capacity of prostate cancer cells was also reduced (Barboro et al., 2015). Migration of hepatoma cells was reduced after PARP inhibition due to altered signaling pathway or proteins involved in migration (Mao et al., 2017). It was recognized in leukocytes that PARP inhibition suppresses certain Rho GTPases and rearrangement of cytoskeletal proteins thereby decreasing adhesion and migration to blood brain barrier (Rom et al., 2016). Proteomic analyses have shown that PARP1 inhibition in cervical cancer cells alters the expression of metastasis-associated genes, particularly those involved in cytoskeletal organization and cell adhesion (Rajawat et al., 2023). In the present study, the phalloidin staining suggested change in cytoskeleton arrangement where we could monitor reduction in cell polarity after PARP inhibition (Figure 2).

To delineate the PARP1 mediated molecular mechanism during breast cancer progression we performed 2DGE and identified key proteins involved in cell-cell adhesion and migration (Figure 3). Here we highlight the role of PARP1 in regulation of cell adhesion and cytoskeletal proteins in triple negative breast cancer. The 2DGE and immunoprecipitation study showed the interaction of vimentin with PARP1. Vimentin is both an EMT and EndoMT marker, and contributes to tumor phenotype and invasiveness (Kidd et al., 2014; McInroy & Maatta, 2007). Moreover, the reduced vimentin levels affect the migration of cells by regulating the intermediary filaments (Mendez et al., 2010). PARP1 has been reported to cause disassembly of vimentin filaments during LPS induced microglia activation (Chen et al., 2024). Interestingly, we have reported the reduction in vimentin level upon PARP inhibition in TNBC cells as confirmed in the western data (Figure 3).

Our 2D-MS proteomics analysis report also shows increased RhoGDIα in MDA-MB-231 cells (Figure 3). The RhoGDIα expression is known to be inversely associated with tumor progression stage where its expression is lost during malignancy and metastasis (Barboro et al., 2015). Another study observed the increase in migration and invasion after downregulation of RhoGDIα in breast cancer cells (Hooshmand et al., 2013). Further, RhoGDIα exhibit multiple functions in regulating members of RhoGTPase family (Dovas & Couchman, 2005; Garcia-Mata et al., 2011), inhibits RhoGTPases, thus reducing the cellular migration (Wollert et al., 2002; Yu et al., 2012). The loss of RhoGDIα signifies increased activity of RhoGTPases and thus enhanced cell motility (Bozza et al., 2015). RhoGDIα is known to regulate RhoGTPases by the Rac/Cdc42/FAK pathway and regulates cell motility and migration. Moreover, RhoGDIα sequesters Rac protein involved in migration (Lawson & Ridley, 2018). RhoGTPases have an important role to play in cellular migration and invasion by reorganizing the cellular actin system (Wang et al., 2014). Suitable distribution of Rho proteins in a cell is essential for their appropriate function and RhoGDIα is known to regulate this localization, thereby regulating RhoGTPase signaling(Lawson & Ridley, 2018). A single report suggests the role of PARP1 in cytoskeletal rearrangement through Rho and Rac proteins in HIV infection, whereby PARP inhibition reduces LTR activity and diminishes RhoGTPases activity (Rom et al., 2015). Hence, monitoring the levels of proteins involved in migration and cytoskeleton associated with RhoGDIα could be of importance in PARP inhibitor therapeutics. Therefore, we performed western analyses for downstream signaling proteins, and found that Rac and Cdc42 expression reduces with PARP inhibition, correlating with increased RhoGDIα (Figure 3). Current study highlights the novel role of PARP1 in regulating cytoskeletal proteins and RhoGTPases machinery through RhoGDIα in breast cancer progression.

PARP1 can regulate the function of several transcription factors, including p53 and NF-kB(Hassa et al., 2003; Hassa et al., 2005; Ko & Ren, 2012; Weaver & Yang, 2013). PARP1 interacts with the transcription factors HIF1α (Elser et al., 2008; Martin-Oliva et al., 2006), Snail1 (Rodriguez et al., 2011) and vimentin (Rodriguez et al., 2013) in various cancer types. Interestingly, in our study we identified transcriptional regulation of vimentin and RhoGDIα by PARP1 as seen from IP and ChIP analyses (Figure 4). RhoGDIα expression studies (Figure 5) supported our data that PARP mediated effects were through modulation of vimentin and RhoGDIα expression. In particular we reported PARP1 interaction with vimentin and RhoGDIα, hence regulating cell migration.

Our animal studies with PARP inhibitor treatment showed significant reduction in tumor weight and volume, as well as metastasis, including bone metastasis (Figure 6). Moreover, immunohistochemistry of clinical samples suggested higher expression of PARP1 and vimentin in triple negative breast cancer patients (Figure 7). Thus, increased PARP1 in breast cancer patients provide an evidence for its role in cancer progression.

The present study is the first experimental evidence that directly implies PARP1 in transcriptional regulation of RhoGDIα. With this work we position PARP1 as one of the key regulators of metastasis in breast cancer. Our findings indicate that PARP inhibitors reduce the metastatic potential of breast cancer cells possibly by regulating the expression of cytoskeletal, cell adhesion and motility proteins, particularly transcriptional regulation of RhoGDIα and vimentin (Figure 8). These proteins then exhibit cumulative effect by regulating the downstream proteins and furthermore, migration and invasion. In summary, we identified novel aspect of PARP1 as promoter of metastasis via transcriptional regulation of vimentin and RhoGDIα in breast cancer cells.

**Figure 8.**
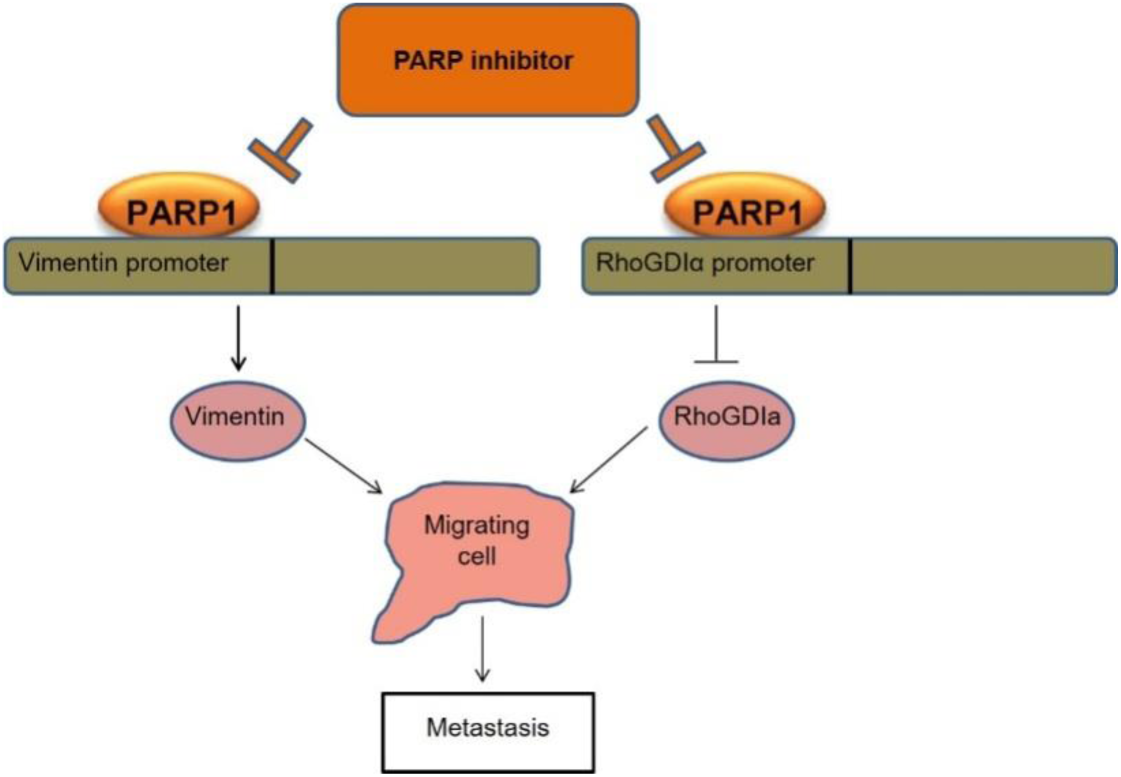
Schematic representation of PARP inhibition on TNBC cells.

## Conclusion

In this study, we propose that PARP1 mediated breast carcinoma progression is gene transcription mediated, where it regulates several steps of pro-metastasis. This study highlights on the efficacy of PARP inhibitors in the setting of metastatic progression of triple negative breast cancer irrespective of BRCA status. Assessing RhoGDIα levels in TNBC patients might be useful to predict sensitivity to PARP inhibitor olaparib. These findings suggest that PARP1 may influence tumor aggressiveness through mechanisms extending beyond DNA repair, underscoring its potential as both a therapeutic target and a biomarker of metastatic progression. Hence, further molecular studies need to be done on large number of patients receiving PARP inhibitor therapy.

## Acknowledgement

This research was supported by the grants from Department of Biotechnology (DBT) to JR (GAP0189) and to DPM (GAP0187). JR thanks DBT for Bio-CARe fellowship (BT/AB/08/01/2008-III) and NS thanks ICMR for SRF fellowship. We thank all the members of the DP Mishra laboratory for helpful discussions. We would like to thank institutional animal house facility for *in vivo* work and SAIF facility for Confocal microscopy.

## Author Contributions

JR conceived the idea, received funding, planned and carried out experiments, analysed the result, designed figures and wrote the manuscript. NS carried out experiments and designed figures. AS, MS and AAJB each carried out one experiment. MS provided the clinical samples. DS reviewed and provided feedback. DPM supervised the findings, aided in data interpretation, edited the manuscript and provided critical feedback.

## References

Ahmad A, Olah G, Herndon DN, & Szabo C (2018). The clinically used PARP inhibitor olaparib improves organ function, suppresses inflammatory responses and accelerates wound healing in a murine model of third-degree burn injury. Br J Pharmacol 175: 232–245.

Bai P, & Virag L (2012). Role of poly(ADP-ribose) polymerases in the regulation of inflammatory processes. FEBS Lett 586: 3771–3777.

Barboro P, Ferrari N, Capaia M, Petretto A, Salvi S, Boccardo S, et al. (2015). Expression of nuclear matrix proteins binding matrix attachment regions in prostate cancer. PARP-1: New player in tumor progression. Int J Cancer 137: 1574–1586.

Beniey M, Hubert A, Haque T, Cotte AK, Bechir N, Zhang X, et al. (2023). Sequential targeting of PARP with carboplatin inhibits primary tumour growth and distant metastasis in triple-negative breast cancer. Br J Cancer 128: 1964–1975.

Bieche I, Pennaneach V, Driouch K, Vacher S, Zaremba T, Susini A, et al. (2013). Variations in the mRNA expression of poly(ADP-ribose) polymerases, poly(ADP-ribose) glycohydrolase and ADP-ribosylhydrolase 3 in breast tumors and impact on clinical outcome. Int J Cancer 133: 2791–2800.

Boulter E, Garcia-Mata R, Guilluy C, Dubash A, Rossi G, Brennwald PJ, et al. (2010). Regulation of Rho GTPase crosstalk, degradation and activity by RhoGDI1. Nat Cell Biol 12: 477–483.

Bozza WP, Zhang Y, Hallett K, Rivera Rosado LA, & Zhang B (2015). RhoGDI deficiency induces constitutive activation of Rho GTPases and COX-2 pathways in association with breast cancer progression. Oncotarget 6: 32723–32736.

Brenner JC, Ateeq B, Li Y, Yocum AK, Cao Q, Asangani IA, et al. (2011). Mechanistic rationale for inhibition of poly(ADP-ribose) polymerase in ETS gene fusion-positive prostate cancer. Cancer Cell 19: 664–678.

Brown JS, O’Carrigan B, Jackson SP, & Yap TA (2017). Targeting DNA Repair in Cancer: Beyond PARP Inhibitors. Cancer Discov 7: 20–37.

Chen K, Li Y, Xu H, Zhang C, Li Z, Wang W, et al. (2017). An analysis of the gene interaction networks identifying the role of PARP1 in metastasis of non-small cell lung cancer. Oncotarget 8: 87263–87275.

Chen R, Xie L, Fan Y, Hua X, & Chung CY (2024). Vesicular translocation of PARP-1 to cytoplasm causes ADP-ribosylation and disassembly of vimentin filaments during microglia activation induced by LPS. Front Cell Neurosci 18: 1363154.

Choi EB, Yang AY, Kim SC, Lee J, Choi JK, Choi C, et al. (2016a). PARP1 enhances lung adenocarcinoma metastasis by novel mechanisms independent of DNA repair. Oncogene 35: 4569–4579.

Choi YE, Meghani K, Brault ME, Leclerc L, He YJ, Day TA, et al. (2016b). Platinum and PARP Inhibitor Resistance Due to Overexpression of MicroRNA-622 in BRCA1-Mutant Ovarian Cancer. Cell Rep 14: 429–439.

Chu S, Xu H, Ferro TJ, & Rivera PX (2007). Poly(ADP-ribose) polymerase-1 regulates vimentin expression in lung cancer cells. Am J Physiol Lung Cell Mol Physiol 293: L1127–1134.

Curtin NJ, & Szabo C (2013). Therapeutic applications of PARP inhibitors: anticancer therapy and beyond. Mol Aspects Med 34: 1217–1256.

Domagala P, Huzarski T, Lubinski J, Gugala K, & Domagala W (2011). PARP-1 expression in breast cancer including BRCA1-associated, triple negative and basal-like tumors: possible implications for PARP-1 inhibitor therapy. Breast Cancer Res Treat 127: 861–869.

Dovas A, & Couchman JR (2005). RhoGDI: multiple functions in the regulation of Rho family GTPase activities. Biochem J 390: 1–9.

Drew Y (2015). The development of PARP inhibitors in ovarian cancer: from bench to bedside. Br J Cancer 113 Suppl 1: S3–9.

Dziaman T, Ludwiczak H, Ciesla JM, Banaszkiewicz Z, Winczura A, Chmielarczyk M, et al. (2014). PARP-1 expression is increased in colon adenoma and carcinoma and correlates with OGG1. PLoS One 9: e115558.

Elser M, Borsig L, Hassa PO, Erener S, Messner S, Valovka T, et al. (2008). Poly(ADP-ribose) polymerase 1 promotes tumor cell survival by coactivating hypoxia-inducible factor-1-dependent gene expression. Mol Cancer Res 6: 282–290.

Faraoni I, Compagnone M, Lavorgna S, Angelini DF, Cencioni MT, Piras E, et al. (2015). BRCA1, PARP1 and gammaH2AX in acute myeloid leukemia: Role as biomarkers of response to the PARP inhibitor olaparib. Biochim Biophys Acta 1852: 462–472.

Farmer H, McCabe N, Lord CJ, Tutt AN, Johnson DA, Richardson TB, et al. (2005). Targeting the DNA repair defect in BRCA mutant cells as a therapeutic strategy. Nature 434: 917–921.

Feng FY, de Bono JS, Rubin MA, & Knudsen KE (2015). Chromatin to Clinic: The Molecular Rationale for PARP1 Inhibitor Function. Mol Cell 58: 925–934.

Frederick MI, Abdesselam D, Clouvel A, Croteau L, & Hassan S (2024). Leveraging PARP-1/2 to Target Distant Metastasis. Int J Mol Sci 25.

Galia A, Calogero AE, Condorelli R, Fraggetta F, La Corte A, Ridolfo F, et al. (2012). PARP-1 protein expression in glioblastoma multiforme. Eur J Histochem 56: e9.

Garcia-Mata R, Boulter E, & Burridge K (2011). The ‘invisible hand’: regulation of RHO GTPases by RHOGDIs. Nat Rev Mol Cell Biol 12: 493–504.

Gil-Kulik P, Dudzinska E, Radzikowska-Buchner E, Wawer J, Jojczuk M, Nogalski A, et al. (2020). Different regulation of PARP1, PARP2, PARP3 and TRPM2 genes expression in acute myeloid leukemia cells. BMC Cancer 20: 435.

Green AR, Caracappa D, Benhasouna AA, Alshareeda A, Nolan CC, Macmillan RD, et al. (2015). Biological and clinical significance of PARP1 protein expression in breast cancer. Breast Cancer Res Treat 149: 353–362.

Guillot C, Favaudon V, Herceg Z, Sagne C, Sauvaigo S, Merle P, et al. (2014). PARP inhibition and the radiosensitizing effects of the PARP inhibitor ABT-888 in in vitro hepatocellular carcinoma models. BMC Cancer 14: 603.

Hassa PO, Buerki C, Lombardi C, Imhof R, & Hottiger MO (2003). Transcriptional coactivation of nuclear factor-kappaB-dependent gene expression by p300 is regulated by poly(ADP)-ribose polymerase-1. J Biol Chem 278: 45145–45153.

Hassa PO, Haenni SS, Buerki C, Meier NI, Lane WS, Owen H, et al. (2005). Acetylation of poly(ADP-ribose) polymerase-1 by p300/CREB-binding protein regulates coactivation of NF-kappaB-dependent transcription. J Biol Chem 280: 40450–40464.

Hooshmand S, Ghaderi A, Yusoff K, Karrupiah T, Rosli R, & Mojtahedi Z (2013). Downregulation of RhoGDIalpha increased migration and invasion of ER (+) MCF7 and ER (-) MDA-MB-231 breast cancer cells. Cell Adh Migr 7: 297–303.

Kaminska K, Szczylik C, Bielecka ZF, Bartnik E, Porta C, Lian F, et al. (2015). The role of the cell-cell interactions in cancer progression. J Cell Mol Med 19: 283–296.

Kidd ME, Shumaker DK, & Ridge KM (2014). The role of vimentin intermediate filaments in the progression of lung cancer. Am J Respir Cell Mol Biol 50: 1–6.

Kim KM, Moon YJ, Park SH, Park HJ, Wang SI, Park HS, et al. (2016). Individual and Combined Expression of DNA Damage Response Molecules PARP1, gammaH2AX, BRCA1, and BRCA2 Predict Shorter Survival of Soft Tissue Sarcoma Patients. PLoS One 11: e0163193.

Ko HL, & Ren EC (2012). Functional Aspects of PARP1 in DNA Repair and Transcription. Biomolecules 2: 524–548.

Korkmaz-Icoz S, Szczesny B, Marcatti M, Li S, Ruppert M, Lasitschka F, et al. (2018). Olaparib protects cardiomyocytes against oxidative stress and improves graft contractility during the early phase after heart transplantation in rats. Br J Pharmacol 175: 246–261.

Kraus WL (2008). Transcriptional control by PARP-1: chromatin modulation, enhancer-binding, coregulation, and insulation. Curr Opin Cell Biol 20: 294–302.

Krukenberg KA, Jiang R, Steen JA, & Mitchison TJ (2014). Basal activity of a PARP1-NuA4 complex varies dramatically across cancer cell lines. Cell Rep 8: 1808–1818.

Kumar M, Jaiswal RK, Prasad R, Yadav SS, Kumar A, Yadava PK, et al. (2021). PARP-1 induces EMT in non-small cell lung carcinoma cells via modulating the transcription factors Smad4, p65 and ZEB1. Life Sci 269: 118994.

Lawson CD, & Ridley AJ (2018). Rho GTPase signaling complexes in cell migration and invasion. J Cell Biol 217: 447–457.

Li W, Wang H, Jin X, & Zhao L (2013). Loss of RhoGDI is a novel independent prognostic factor in hepatocellular carcinoma. Int J Clin Exp Pathol 6: 2535–2541.

Lin K, Zhao Y, Tang Y, Chen Y, Lin M, & He L (2024). Collagen I-induced VCAN/ERK signaling and PARP1/ZEB1-mediated metastasis facilitate OSBPL2 defect to promote colorectal cancer progression. Cell Death Dis 15: 85.

Mao X, Du S, Yang Z, Zhang L, Peng X, Jiang N, et al. (2017). Inhibitors of PARP-1 exert inhibitory effects on the biological characteristics of hepatocellular carcinoma cells in vitro. Mol Med Rep 16: 208–214.

Marques M, Beauchamp MC, Fleury H, Laskov I, Qiang S, Pelmus M, et al. (2015). Chemotherapy reduces PARP1 in cancers of the ovary: implications for future clinical trials involving PARP inhibitors. BMC Med 13: 217.

Martin-Oliva D, Aguilar-Quesada R, O’Valle F, Munoz-Gamez JA, Martinez-Romero R, Garcia Del Moral R, et al. (2006). Inhibition of poly(ADP-ribose) polymerase modulates tumor-related gene expression, including hypoxia-inducible factor-1 activation, during skin carcinogenesis. Cancer Res 66: 5744–5756.

McGrail DJ, Lin CC, Garnett J, Liu Q, Mo W, Dai H, et al. (2017). Improved prediction of PARP inhibitor response and identification of synergizing agents through use of a novel gene expression signature generation algorithm. NPJ Syst Biol Appl 3: 8.

McInroy L, & Maatta A (2007). Down-regulation of vimentin expression inhibits carcinoma cell migration and adhesion. Biochem Biophys Res Commun 360: 109–114.

Mendez MG, Kojima S, & Goldman RD (2010). Vimentin induces changes in cell shape, motility, and adhesion during the epithelial to mesenchymal transition. FASEB J 24: 1838–1851.

Pashaiefar H, Yaghmaie M, Tavakkoly-Bazzaz J, Ghaffari SH, Alimoghaddam K, Momeny M, et al. (2018). PARP-1 Overexpression as an Independent Prognostic Factor in Adult Non-M3 Acute Myeloid Leukemia. Genet Test Mol Biomarkers 22: 343–349.

Rajawat J, Awasthi P, & Banerjee M (2023). PARP inhibitor olaparib induced differential protein expression in cervical cancer cells. J Proteomics 275: 104823.

Rajawat J, Shukla N, & Mishra DP (2017). Therapeutic Targeting of Poly(ADP-Ribose) Polymerase-1 (PARP1) in Cancer: Current Developments, Therapeutic Strategies, and Future Opportunities. Med Res Rev 37: 1461–1491.

Reyes SB, Narayanan AS, Lee HS, Tchaicha JH, Aldape KD, Lang FF, et al. (2013). alphavbeta8 integrin interacts with RhoGDI1 to regulate Rac1 and Cdc42 activation and drive glioblastoma cell invasion. Mol Biol Cell 24: 474–482.

Rodriguez MI, Gonzalez-Flores A, Dantzer F, Collard J, de Herreros AG, & Oliver FJ (2011). Poly(ADP-ribose)-dependent regulation of Snail1 protein stability. Oncogene 30: 4365–4372.

Rodriguez MI, Peralta-Leal A, O’Valle F, Rodriguez-Vargas JM, Gonzalez-Flores A, Majuelos-Melguizo J, et al. (2013). PARP-1 regulates metastatic melanoma through modulation of vimentin-induced malignant transformation. PLoS Genet 9: e1003531.

Rom S, Reichenbach NL, Dykstra H, & Persidsky Y (2015). The dual action of poly(ADP-ribose) polymerase-1 (PARP-1) inhibition in HIV-1 infection: HIV-1 LTR inhibition and diminution in Rho GTPase activity. Front Microbiol 6: 878.

Rom S, Zuluaga-Ramirez V, Reichenbach NL, Dykstra H, Gajghate S, Pacher P, et al. (2016). PARP inhibition in leukocytes diminishes inflammation via effects on integrins/cytoskeleton and protects the blood-brain barrier. J Neuroinflammation 13: 254.

Schacke M, Kumar J, Colwell N, Hermanson K, Folle GA, Nechaev S, et al. (2019). PARP-1/2 Inhibitor Olaparib Prevents or Partially Reverts EMT Induced by TGF-beta in NMuMG Cells. Int J Mol Sci 20.

Schiewer MJ, & Knudsen KE (2014). Transcriptional roles of PARP1 in cancer. Mol Cancer Res 12: 1069–1080.

Singh S, Sadacharan S, Su S, Belldegrun A, Persad S, & Singh G (2003). Overexpression of vimentin: role in the invasive phenotype in an androgen-independent model of prostate cancer. Cancer Res 63: 2306–2311.

Siraj AK, Pratheeshkumar P, Parvathareddy SK, Divya SP, Al-Dayel F, Tulbah A, et al. (2018). Overexpression of PARP is an independent prognostic marker for poor survival in Middle Eastern breast cancer and its inhibition can be enhanced with embelin co-treatment. Oncotarget 9: 37319–37332.

Stanley J, Klepczyk L, Keene K, Wei S, Li Y, Forero A, et al. (2015). PARP1 and phospho-p65 protein expression is increased in human HER2-positive breast cancers. Breast Cancer Res Treat 150: 569–579.

Swindall AF, Stanley JA, & Yang ES (2013). PARP-1: Friend or Foe of DNA Damage and Repair in Tumorigenesis? Cancers (Basel) 5: 943–958.

Thakur R, Trivedi R, Rastogi N, Singh M, & Mishra DP (2015). Inhibition of STAT3, FAK and Src mediated signaling reduces cancer stem cell load, tumorigenic potential and metastasis in breast cancer. Sci Rep 5: 10194.

van Zijl F, Krupitza G, & Mikulits W (2011). Initial steps of metastasis: cell invasion and endothelial transmigration. Mutat Res 728: 23–34.

Vasko V, Espinosa AV, Scouten W, He H, Auer H, Liyanarachchi S, et al. (2007). Gene expression and functional evidence of epithelial-to-mesenchymal transition in papillary thyroid carcinoma invasion. Proc Natl Acad Sci U S A 104: 2803–2808.

Wang MJ, Artemenko Y, Cai WJ, Iglesias PA, & Devreotes PN (2014). The directional response of chemotactic cells depends on a balance between cytoskeletal architecture and the external gradient. Cell Rep 9: 1110–1121.

Weaver AN, & Yang ES (2013). Beyond DNA Repair: Additional Functions of PARP-1 in Cancer. Front Oncol 3: 290.

Wei H, & Yu X (2016). Functions of PARylation in DNA Damage Repair Pathways. Genomics Proteomics Bioinformatics 14: 131–139.

Wollert T, DePina AS, Thompson RF, & Langford GM (2002). GTPase rho is involved in myosin-II-mediated contraction of pseudo-contractile rings and transport of vesicles in extracts of clam oocytes. Biol Bull 203: 208–210.

Wu F, Hu P, Li D, Hu Y, Qi Y, Yin B, et al. (2016). RhoGDIalpha suppresses self-renewal and tumorigenesis of glioma stem cells. Oncotarget 7: 61619–61629.

Xiao Y, Lin VY, Ke S, Lin GE, Lin FT, & Lin WC (2014). 14-3-3tau promotes breast cancer invasion and metastasis by inhibiting RhoGDIalpha. Mol Cell Biol 34: 2635–2649.

Yap TA, Kristeleit R, Michalarea V, Pettitt SJ, Lim JSJ, Carreira S, et al. (2020). Phase I trial of the poly(ADP-ribose) polymerase (PARP) inhibitor olaparib and AKT inhibitor capivasertib in patients with BRCA1/2 and non-BRCA1/2 mutant cancers. Cancer Discov.

Yu J, Zhang D, Liu J, Li J, Yu Y, Wu XR, et al. (2012). RhoGDI SUMOylation at Lys-138 increases its binding activity to Rho GTPase and its inhibiting cancer cell motility. J Biol Chem 287: 13752–13760.

Zhang B, Zhang Y, Dagher MC, & Shacter E (2005). Rho GDP dissociation inhibitor protects cancer cells against drug-induced apoptosis. Cancer Res 65: 6054–6062.

Zhu J, Li Y, Chen C, Ma J, Sun W, Tian Z, et al. (2017). NF-kappaB p65 Overexpression Promotes Bladder Cancer Cell Migration via FBW7-Mediated Degradation of RhoGDIalpha Protein. Neoplasia 19: 672–683.

Zhu Y, Tummala R, Liu C, Nadiminty N, Lou W, Evans CP, et al. (2012). RhoGDIalpha suppresses growth and survival of prostate cancer cells. Prostate 72: 392–398.

