## Supplementary figures and images for "PARP1 inhibition regulates tumor progression through modulation of RhoGDIα and vimentin in triple negative breast cancer"

### Supplementary Table 1

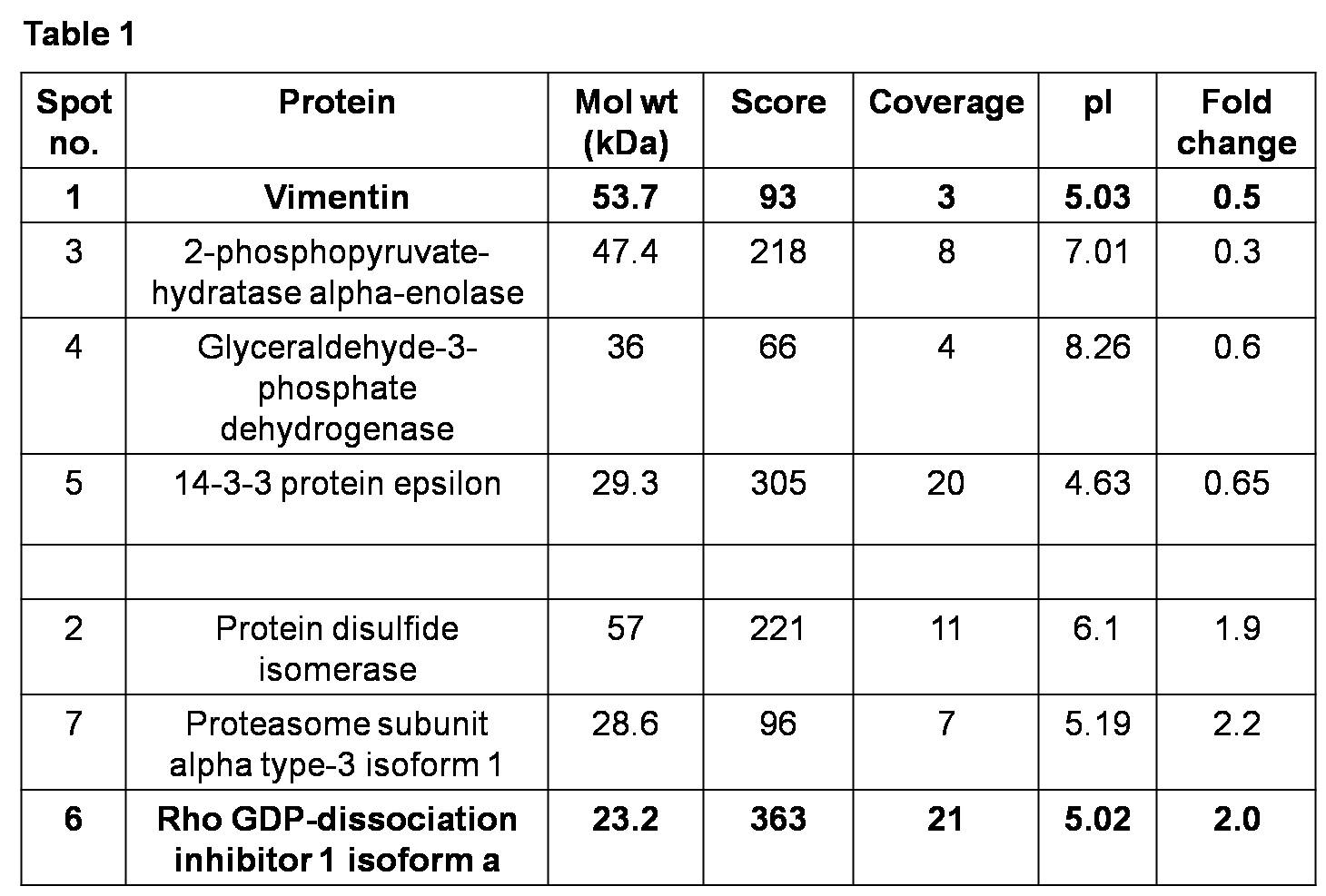
